# Cell-specific wiring routes information flow through hippocampal CA3

**DOI:** 10.1101/2024.06.24.600436

**Authors:** Jake F. Watson, Victor Vargas-Barroso, Peter Jonas

## Abstract

The hippocampus is dogmatically described as a trisynaptic circuit. Dentate gyrus granule cells, CA3 pyramidal neurons (PNs), and CA1 PNs are serially connected, forming a circuit that critically enables memory storage in the brain. However, fundamental aspects of hippocampal function go beyond this simplistic ‘trisynaptic’ definition. CA3 PNs connect not only to CA1, but also establish the largest autoassociative network in the brain. In addition, CA3 PNs are not uniform, differing in their morphology, intrinsic properties, and even the extent of granule cell input. Understanding how these different subtypes of CA3 PNs are embedded in the hippocampal network is essential for our quest to understand learning and memory. Here, we performed simultaneous multi-cellular patch-clamp recordings from up to eight CA3 PNs in acute mouse hippocampal slices, testing 3114 possible connections between identified cells. Combined with post-hoc morphological analysis, this allowed full characterization of neuronal heterogeneity in functioning microcircuits. We demonstrate that CA3 PNs can be divided into distinct ‘deep’ and ‘superficial’ subclasses, with altered input-output balance. While both subtypes formed recurrent connectivity within classes, connectivity between subtypes was surprisingly asymmetric. Recurrent connectivity was abundant from superficial to deep, but almost absent from deep to superficial PNs, thereby splitting CA3 into parallel recurrent networks which will allow more complex information processing. Finally, we observed innervation of PN subclasses by distinct interneurons, a potential mechanism to gate information flow through CA3 sublayers. Together, our data present a major revision to the classical ‘trisynaptic’ view of the hippocampus, bringing us closer to understanding its complex action in information storage.

## Introduction

The hippocampus is fundamental for learning and memory. Famously, patients with hippocampal removal have a stark inability to lay down long-term memories (Squire et al., 2004; Squire, 2009). Therefore, neuronal activity flowing through hippocampal microcircuits is in some way processed and directed to allow information storage. This brain area is classically depicted as a trisynaptic circuit, in which granule cells (GCs), CA3 pyramidal neurons, and CA1 pyramidal neurons (PNs) are connected in series (Andersen et al., 1971; Amaral and Witter, 1989). While this axis forms the backbone of hippocampal information flow, it omits wiring that is also crucial for circuit processing. Most importantly, CA3 PNs are connected to one another, establishing an largest autoassociative network in the brain (Witter, 2007). In addition, not only GCs, but also both CA3 and CA1 PNs receive direct projections from entorhinal layers (Yeckel and Berger, 1990; Ben-Simon et al., 2022), and CA2 neurons add a further synapse in the chain (Dudek et al., 2016). Therefore, the routemap for information flow through even the intensively studied hippocampus is more complex than previously anticipated.

Increased complexity does not stop at wiring. While traditionally assumed to be uniform, substantial cellular heterogeneity of hippocampal PN populations is increasingly emerging. In CA1, superficial calbindin^+^ cells and deep calbindin^−^ neurons can be distinguished (Lorente de Nó, 1934; Celio, 1990), and have different layer densities, genetics, and wiring (Slomianka et al., 2011; Lee et al., 2014), resulting in specialized contributions to population coding (Valero et al., 2015). However, in hippocampal CA3, the organizational principles are less clear. Heterogeneity of morphology and firing properties have been described on both deep-superficial (Fitch et al., 1989; Bilkey and Schwartzkroin, 1990) and proximal-distal hippocampal axes (Kowalski et al., 2016; Sun et al., 2017). Recent work suggested distinct CA3 PN classes based on burst-firing properties and the presence (thorny) or absence (athorny) of mossy fiber input (Hunt et al., 2018). However others have described burst firing PNs with less strict anatomical features (Raus Balind et al., 2019; Magó et al., 2021; Balleza-Tapia et al., 2022). Whether these observations represent gradients of heterogeneity amongst CA3 PNs, or how they map onto gene-expression based clusters of PNs remains to be clarified (Thompson et al., 2008; Cembrowski and Spruston, 2019; Yao et al., 2021). In addition, for hippocampal processing, not only the properties, but the synaptic wiring of these subtypes will critically define the processing capabilities of the hippocampal circuit. CA3-CA3 recurrent synapses are putative sites for information storage, allowing association between incoming activity streams (Rolls, 2018). Currently, subtype-specific recurrent wiring rules remain unknown. To address these questions, we combined multicellular patch clamp-based circuit mapping with post-hoc morphological analysis. This approach allowed us to determine synaptic connectivity of rigorously identified, physiologically and morphologically characterized CA3 pyramidal cells. We found that superficial and deep CA3 PNs differed in physiological properties, morphological characteristics, and crucially, synaptic interconnectivity. These properties lead to a bifurcation of the transhippocampal information stream, initiating parallel signal processing in the center of the hippocampal circuit. Preliminary accounts of the work were previously published in abstract form (Watson et al., 2022).

## Results

### Different CA3 PN subclasses coexist on the deep-superficial axis

To determine the connectivity properties in the CA3 recurrent circuitry, we performed multicellular patch-clamp based circuit mapping in acute slices of mouse hippocampus. In total, we recorded 928 CA3 PNs. Coupled with post-hoc visualization using 3,3’-diaminobenzidine (DAB) as chromogen, this approach allows full functional and morphological analysis of cellular heterogeneity at the microcircuit level (**Fig. 1A**). A hallmark feature of CA3 PNs is the presence of complex ‘thorny excrescences’ on primary dendrites, the postsynaptic sites of GC input (Vandael and Jonas, 2024), yet a subpopulation of neurons lacking these structures has been reported (athorny neurons) (Hunt et al., 2018). Visualization of our recorded cells confirmed multiple examples of such neurons (**Fig. 1B**), which, as characterized (Hunt et al., 2018; Linaro et al., 2022) also displayed ‘burst-like’ activity patterns on current injection and had a deep somatic location in *stratum pyramidale* (**Fig. 1C**). These neurons have a stereotypical dendritic arborisation pattern, where the primary dendrite crosses *stratum pyramidale* before branching. In addition to these cells, we also observed a substantial proportion of neurons with a similar morphology, somatic location, and burst-firing profile to ‘athorny’ neurons, which also had sparse thorny excrescences (**Fig. 1D**).

**Fig. 1.**
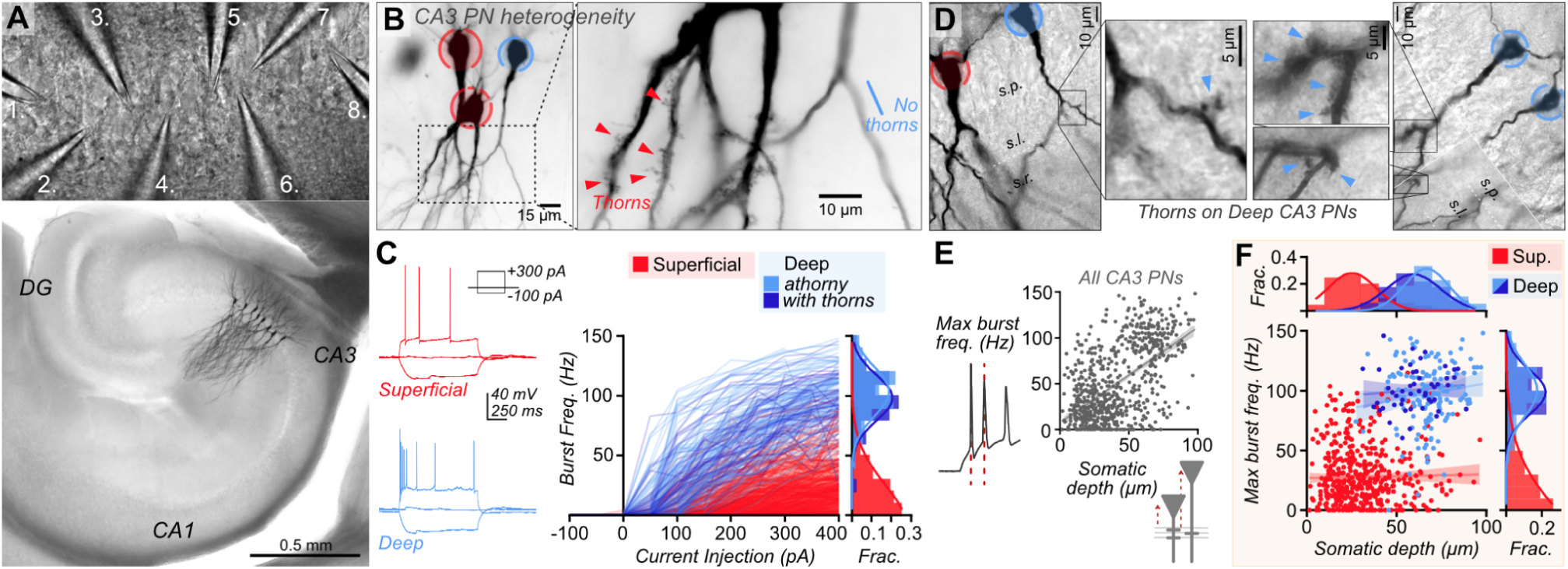
| CA3 PNs form superficial and deep subclasses. **A** Infrared-DIC view of microelectrode pipettes (numbered) during multicellular recording (upper), and DAB-stained hippocampal slice after octuple recording in CA3 (lower). **B** Light micrographs of CA3 PNs show cells with (red) or without (blue) thorny excrescences. **C** Analysis of AP phenotypes evoked by long current pulses (representative traces, left), showed distinct trajectories of initial burst frequency upon increasing current injection between PN classes (thorny/superficial and athorny/deep). A population of neurons with overall athorny morphology and somatic location, but evident thorny excrescences had a similar firing profile to athorny neurons. Burst frequency at 400 pA injection is also presented as a relative fraction histogram, fitted with Gaussian curves (right) **D** Example images of putative ‘deep’ PNs displaying equivalent morphology to athorny neurons, but with sparse thorny excrescences (arrows). s.p. = *stratum pyramidale*, s.l. = *stratum lucidum*, s.r. = *stratum radiatum*, layer boundaries denoted by dashed lines. A composite z-plane image (right) is presented for visualization of multiple structures. Boundaries between images are depicted with hard lines. **E** Scatter plots of maximal burst frequency upon current injection and somatic depth (distance from *stratum lucidum*) for all CA3 PNs shows distinct clusters and overall positive correlation (linear regression: slope = 1.04 Hz μm^-1^, R^2^ = 0.33, p < 0.0001). **F** Morphological assignment of PNs shows almost complete separation of PNs into the two clusters. Morphologically ‘athorny’ neurons with evident thorns (dark blue) populate the ‘deep’ cluster, demonstrating that athorny neurons form part of a larger subclass of ‘deep CA3 PNs’. No significant correlation between maximal burst frequency and somatic depth was observed within clusters, confirming that PNs form discrete subclasses, rather than one heterogeneous gradient (linear regressions; superficial: slope 0.03 Hz μm^-1^, R^2^ = 0.00, p = 0.63; deep (athorny): slope 0.28 Hz μm^-1^, R^2^ = 0.02, p = 0.10; deep (with thorns): slope 0.14 Hz μm^-1^, R^2^ = 0.01, p = 0.46). Relative fractional histograms for each axis are depicted with Gaussian fits.

We therefore sought to determine if putative ‘athorny’ neurons were indeed a discrete subclass of PNs, or represent the end of a heterogeneous gradient of neurons along the deep-superficial axis, with differing mossy fiber input and firing patterns. We analyzed the firing properties of all recorded CA3 PNs with sufficient recording and morphological quality (see Methods), plotting the maximal burst frequency upon current injection against somatic depth (distance from *stratum lucidum*). Amongst all PNs, we observed a positive correlation between burst frequency and somatic depth, however neurons were clustered into two clearly visible populations (**Fig. 1E**). Mapping morphological identity onto these clusters showed almost perfect segregation between neurons with ‘athorny’ morphology and classical CA3 PNs (**Fig. 1F**). However, athorny neurons clustered with sparsely thorny burst-firing neurons. Therefore, while CA3 PNs can be dissected into discrete subclasses, athorny neurons are a subpopulation of a broader subclass of PNs for which ‘lack of thorny excrescences’ is not a defining criteria. These conclusions are consistent with other recent analysis in dorsal CA3 (Magó et al., 2021). For these reasons, a more appropriate terminology for the two CA3 PN subpopulations would be ‘deep’ and ‘superficial’ neurons, which we use henceforth.

To confirm that deep and superficial neurons were indeed distinct subclasses, we plotted linear regressions within morphologically identified clusters. If neurons were a continuous gradient, we would expect the linear relationship observed across all neurons to be maintained within subclusters. However, this was not the case. Positive correlation between burst frequency and somatic depth was abolished within subclasses (**Fig. 1F**). Therefore CA3 PNs can be separated into distinct subclasses based on firing properties and somatic location. Linear discriminant analysis (LDA) could reliably segregate subclasses, with 94% agreement to morphological classification alone. This served as a basis to assign all recorded neurons to either deep or superficial identity (**Supplementary Fig. 1**, see Materials and Methods for complete description), resulting in 531 putative superficial and 233 deep PNs. A minor subset of neurons had morphology substantially deviating from their assigned cluster, and were excluded from further analysis (putative superficial thorn-lacking neurons (n = 2), and extensively thorny, deep bursting neurons with multiple primary apical dendrites (n = 3)). It is important to note that electrophysiological recordings are not an unbiased means to quantify the exact proportion of cell types. Due to deliberate targeting of deep PNs for characterisation, our dataset will overrepresent the proportion of this subclass, which is more likely to make up around 10–20% of all CA3 PNs (Hunt et al., 2018).

### Asymmetric synaptic connectivity between deep and superficial CA3 PNs

Next, we analyzed the local synaptic connectivity between PNs in the CA3 recurrent collateral network, focusing on source– and target cell-specific differences in synaptic connectivity (**Fig. 2A-C**). After PN subclassification, our dataset assessed 3114 putative connections between identified CA3 PNs, observing 85 monosynaptic connections. Due to slicing, this is likely an underestimate of synaptic connectivity in the intact brain, which we have previously estimated to be at least 1.4-fold higher than recorded in slices without further corrections (Watson et al., 2024). Connectivity was no different between sexes or hemispheres (**Supplementary Fig. 2A**), therefore all data were pooled.

**Fig. 2.**
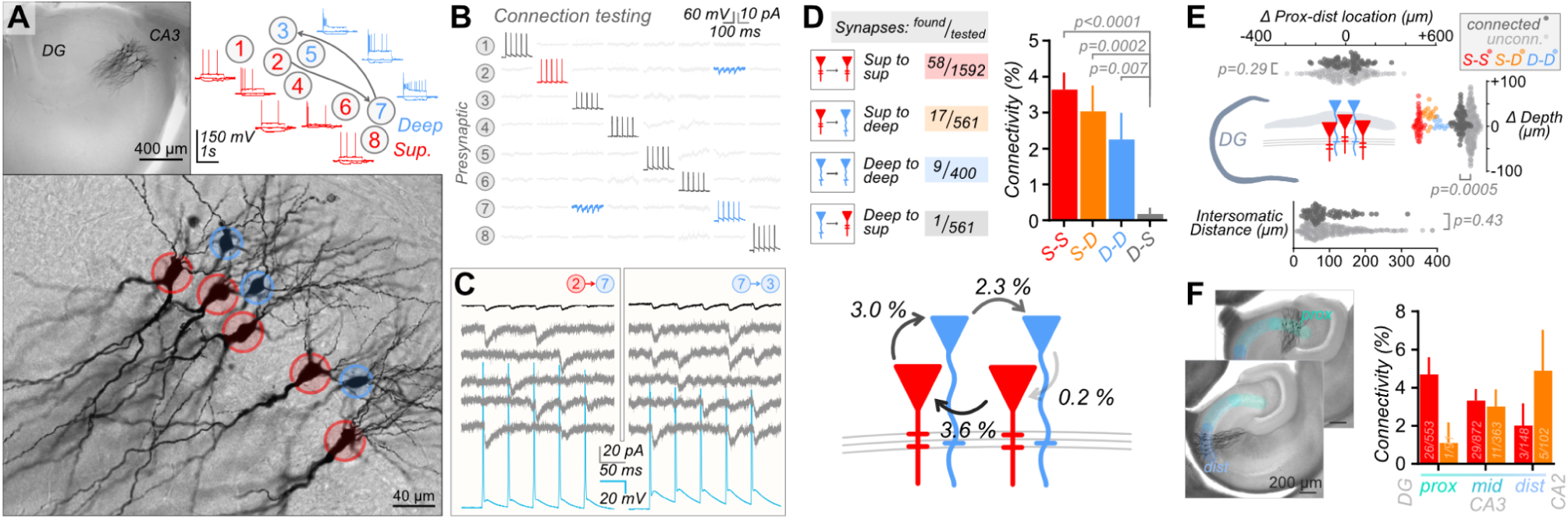
| Deep and superficial subclasses have asymmetric connectivity in the CA3 recurrent network. **A** Example octuple recording from CA3, including both deep (blue) and superficial (red) subclasses. Micrographs at low (upper) and high (lower) resolution depict neuronal morphologies, with cell identity labeled (color). Identified synaptic connectivity (arrows) is schematically depicted, alongside firing profiles for each neuron (−100, 0 and +400 pA injection traces displayed). **B** Voltage-clamp connectivity testing matrix for cells recorded in **A**, with monosynaptic connections presented as bold, and traces involved in connections colored by cell-type. **C** Synaptic traces from **B**. Presynaptic APs (cyan) are followed at short latency by postsynaptic EPSCs. Both postsynaptic individual traces (grey) and average response (black) are presented. **D** Pooled connectivity measures from all recordings demonstrate asymmetric connectivity in the CA3 recurrent network. Connections from superficial PNs to both superficial and deep PNs are evident, yet deep PNs connect almost exclusively to other deep PNs. Error bars present SD estimated from a binomial distribution. Connectivity %, measured ± SD; S-S: 3.64 ± 0.47%; S-D: 3.03 ± 0.72%; D-D: 2.25 ± 0.74%; D-S: 0.18 ± 0.18%; Fisher’s exact test, p < 0.0001, p values of pairwise comparisons after Benjamini Hochberg correction are labeled. **E** No bias for directional connectivity on the proximal-distal axis was observed (upper; + values denote postsynaptic soma more distal (towards CA2/1) than presynaptic soma; unconnected, −0.1 ± 2.1 μm, n = 2910; connected, 12.3 ± 11.4 μm, n = 81, Mann-Whitney test, p = 0.29). In addition, no intersomatic distance dependence to connectivity was observed (all PN subclass intersomatic distance, lower: unconnected, 97.7 ± 1.1 μm, n = 2936; connected, 92.1 ± 5.9 μm, n = 83; Mann-Whitney test, p = 0.43). However, overall CA3 PN connectivity was biased for connectivity towards deeper PNs (+ values denote deeper postsynaptic soma. All PN subclasses, right: unconnected, −0.3 ± 0.6 μm, n = 2996; connected, 10.5 ± 2.4 μm, n = 85, Mann-Whitney test, p = 0.0005), likely in part due to asymmetric connectivity. Individual subclass distances are also plotted (color). **F** Micrographs of example recordings from proximal (upper) or distal (lower) CA3. Breakdown of connectivity into proximal, mid or distal CA3 for S-S and S-D connections (Connectivity %, measured ± SD; Proximal – S-S: 4.7 ± 0.9%, S-D: 1.1 ± 1.1%; Mid – S-S: 3.3 ± 0.6%, S-D: 3.0 ± 0.9%; Distal – S-S: 2.0 ± 1.2%, S-D: 4.9 ± 2.1%; Fisher’s exact test, p = 0.37).

Surprisingly, we discovered highly asymmetric wiring rules (**Fig. 2D**): while superficial CA3 cells provided input to both PN subclasses (to sup: 58/1592 and to deep: 17/561 identified connections), deep cells showed prominent connectivity only to other deep PNs (9/400 connections). Only one CA3-CA3 connection was observed from deep to superficial CA3 PNs (1/561 connections). Our results suggest that the CA3 PN recurrent network is not a simple all-to-all recurrent network, but subclass heterogeneity generates more complex activity flow. Superficial CA3 PNs appear to form the ‘textbook-style’ CA3 recurrent circuit, yet deep CA3 PNs receive the output of this network, forming a separate recurrent network that sends little projection back to superficial CA3 at the local level.

CA3 is thought to form a broad recurrent network across the hippocampal structure. In line with this, we observed no bias for directional connectivity on the proximal-distal axis, with connections between neurons occurring both in the direction of the DG, and CA2/1 (**Fig. 2E, Supplementary Fig. 2B**). No intersomatic distance dependence of connectivity was observed either, in line with other reports in the CA3 of rats, mice, and humans (Guzman et al., 2016; Sammons et al., 2024; Watson et al., 2024) (**Fig. 2E, Supplementary Fig. 2C**). We did however observe a bias for connectivity towards deeper neurons (**Fig. 2E**), which is predominantly generated by the asymmetric connectivity between PN subclasses, but is also observed trendwise within individual subclass datasets (**Supplementary Fig. 2D**). Previous work demonstrated clear changes in the properties of CA3 PNs on the transverse (proximal to distal) axis (Kowalski et al., 2016; Sun et al., 2017). We could confirm the tendency for higher firing in proximal CA3 from our dataset (**Supplementary Fig. 3A, B**). This region also appeared to contain the fewest recorded ‘deep’ CA3 PNs, correlating with the presence of the infrapyramidal blade of the mossy fiber tract (**Supplementary Fig. 3C**). Calculation of connectivity on this axis also showed an apparent bias in the targeting of superficial PN axons, from other superficial neurons in proximal CA3, to more deep PNs in distal CA3 (**Fig. 2F**). However, more extensive analysis is required to confirm these tendencies.

In summary, we demonstrate subtype-specific differences in connectivity between superficial and deep CA3 PNs. Thus, the hippocampal CA3 region is not simply part of the trisynaptic circuit, but rather forms a two-layered circuit in which incoming information is processed in two separate layers of the recurrent network.

### Similar synaptic properties of unitary connections between CA3 PN subclasses

We next studied the synaptic properties of unitary connections between CA3 PNs in the recurrent system. Excitatory postsynaptic current (EPSC) recordings displayed no subclass-dependent differences in potency, success rate, or latency of connections between CA3 PNs (**Fig. 3A**). Synaptic kinetics were also similar across subclasses (**Fig. 3B**). During 20-Hz trains, we observed subtly depressing average responses, which were driven by decreasing success rates of events with similar potency (**Fig. 3C**). Again, these properties appeared similar across PN subtypes. Together, it appears that while CA3 recurrent synapses have heterogeneous properties, they are used similarly across PN subclasses.

**Fig. 3.**
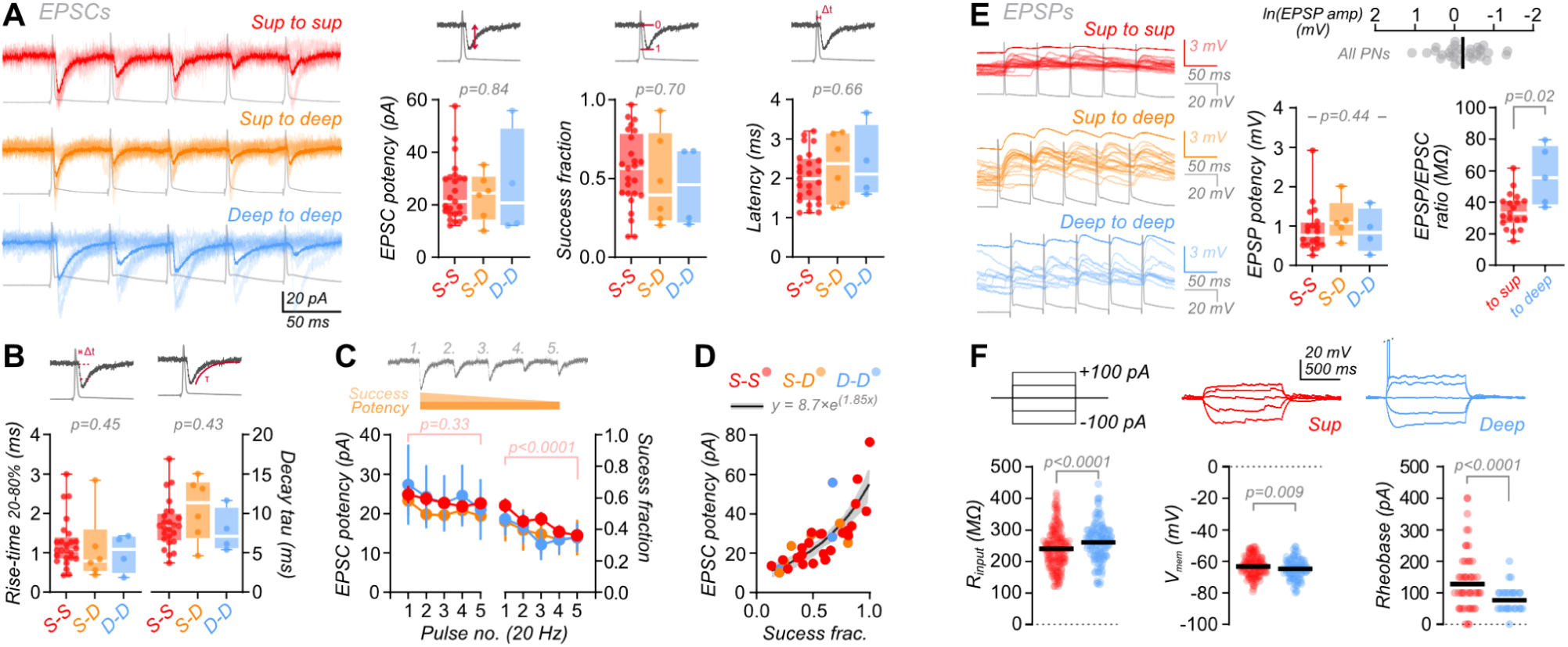
| Recurrent synaptic properties are independent of CA3 PN subclass. **A** EPSCs of recurrent synapses between recorded CA3 PNs. Synaptic currents show no subclass-specific properties, with similar potencies (S-S: 24.9 ± 2.2 pA, n = 25; S-D: 23.3 ± 3.7 pA, n = 6; D-D: 27.3 ± 10.2 pA, n = 4. Kruskal-Wallis test, p = 0.84), success fraction (S-S: 0.55 ± 0.05, n = 25; S-D: 0.49 ± 0.12, n = 6; D-D: 0.45 ± 0.13, n = 4; Kruskal-Wallis test, p = 0.70), and latencies (S-S: 2.03 ± 0.13 ms, n = 25; S-D: 2.28 ± 0.35 ms, n = 6; D-D: 2.37 ± 0.47 ms, n = 4; Kruskal-Wallis test, p = 0.66). **B** EPSC kinetics also remain unchanged between subclasses (Rise time (20–80%) – S-S: 1.24 ± 0.12 ms, n = 25; S-D: 1.10 ± 0.36; D-D: 1.00 ± 0.24, n = 4; Kruskal-Wallis test, p = 0.45. Decay tau – S-S: 8.6 ± 0.6 ms, n = 25; S-D: 10.6 ± 1.6, n = 6; D-D: 7.8 ± 1.4, n = 4; Kruskal-Wallis test, p = 0.43). **C** Responses across 20-Hz train stimulation are produced by events with similar potency (left), but decreasing fraction of successes (right) (EPSC potency, 2-way ANOVA with Tukey’s multiple comparison tests: S-S Pulse 1 vs. 5, p = 0.33. Success fraction, S-S Pulse 1 vs. 5, p < 0.0001). **D** The relationship between synaptic potency and success fraction is independent of PN subclass, but shows a striking exponential relationship (Exponential fit of all datapoints, y=c*exp(kx), R^2^ = 0.65. Fit and 95% confidence intervals plotted as line and shading respectively). **E** Overall, EPSPs follow a log-normal distribution, as demonstrated by plotting the natural log of amplitudes (upper; line at mean (−0.22 mV); Shapiro-Wilk normality test, W = 0.98, p = 0.90). EPSP amplitudes show no significant difference between PN subclasses (S-S: 0.92 ± 0.13 mV, n = 20; S-D: 1.15 ± 0.24 mV, n = 5; D-D: 0.88 ± 0.28 mV, n = 4; Kruskal-Wallis test, p = 0.44), but the EPSP-to-EPSC ratio of CA3 synapses onto deep PNs is greater than onto superficial PNs (CA3 PN synapses onto sup: 34.7 ± 2.6 MΩ, n = 19; onto deep PNs, 56.9 ± 8.4 MΩ, n = 5; Mann-Whitney test, p = 0.02). **F** Intrinsic properties may explain this effect, with a higher input resistance at deep than superficial PNs (Example traces have spikes truncated for presentation (dashed line). Plots have line at mean. R_input_ – superficial PNs (red): 239.9 ± 3.2 MΩ, n = 354; deep (blue): 260.7 ± 4.5 MΩ, n = 171; Mann-Whitney test, p < 0.0001). Resting membrane potential is subtly lower in deep PNs, (line at mean, sup: −63.2 ± 0.3 mV, n = 354; deep: −64.7 ± 0.47 mV, n = 171; Welch’s t test, p = 0.009). The lower rheobase (current required for firing) in deep PNs is also a likely consequence of membrane properties (line at mean. sup: 127.7 ± 4.1 pA, n = 354; deep: 77.2 ± 2.6 pA, n = 171; Mann-Whitney test, p < 0.0001), producing a CA3 PN built to respond more rapidly to input currents.

When analyzing these properties further, we observed a striking relationship between EPSC potency and success rate, following an exponential function (**Fig. 3D**). Weaker synapses, with lower synaptic potency not only produce a smaller synaptic depolarisation, but are also the most unreliable. In contrast, more reliable synapses are also exponentially more potent, producing an ∼80-fold disparity in the overall transmission strength between the weakest and strongest synapses. This relationship is consistent with a substantial variability in release probability or number of functional release sites across CA3-CA3 synapses. This dataset of synaptic properties across the CA3 recurrent circuit reflects the result of ongoing synaptic plasticity or engram storage. Therefore there is clearly a significant presynaptic component to long-term storage of synaptic strength at this connection, which works in tandem with postsynaptic mechanisms.

We next examined excitatory postsynaptic potential (EPSP) properties. As previously observed in multiple species, (Guzman et al., 2016; Miles and Wong, 1986; Sammons et al., 2024; Watson et al., 2024), CA3-CA3 synapses are relatively weak, with mean unitary depolarisations across all subclasses of just 0.94 ± 0.10 mV (**Fig. 3E**). Therefore substantial synaptic integration or summation will be required for spike generation by recurrent activity, and associational network function. We observed no difference in EPSP potency between connection subtypes, however comparing the ratio of EPSP and EPSC at the same synapses revealed a higher EPSP output for CA3 synapses onto deep PNs than superficial PNs (**Fig. 3E**). To probe the mechanisms underlying this difference, we measured the input resistance of the postsynaptic superficial and deep neurons. As previously shown, input resistance was significantly higher in deep than in superficial neurons (Balleza-Tapia et al., 2022; Hunt et al., 2018; Linaro et al., 2022) (**Fig. 3F**, 240 ± 3 MΩ vs. 261 ± 4 MΩ, p < 0.0001). This membrane property also likely underlies the lower rheobase current observed for deep PNs (**Fig. 3F**), a further example of enhanced output properties for deep PNs. Overall, we observe no synapse specific properties of CA3 subclasses, suggesting that CA3 recurrent synapses form a single population. However, how these synapses are used for their computational purpose will be determined by the cell subclass they belong to, both through cell-specific input/output properties, and the asymmetric wiring rules that we have uncovered.

Further subclassification of non-bursting PNs into weak and strongly adapting cells has been performed previously (Hemond et al., 2008; Balleza-Tapia et al., 2022). Our recordings reflect these findings within the superficial subclass (**Supplementary Fig. 4**). However, we observed little spatial, functional, or connective differences between these cells. Therefore the significance of these differences for CA3 function remains to be clarified.

### Correlated spontaneous activity suggests differential inhibitory innervation of CA3 PN subclasses

Our recordings allow analysis not only of evoked synaptic transmission between cells, but also of spontaneous input from unrecorded cells elsewhere in the slice, giving a measure of the input levels to recorded cells. The frequency of spontaneous PSCs (sPSCs) was significantly higher on superficial than deep PNs (5.4 ± 0.5 vs. 3.0 ± 0.4 Hz, **Fig. 4A**). GABAzine application (10 μM) blocked a similar frequency of events between subclasses (sup: 2.2, deep: 2.0 Hz), leaving a far lower frequency of sEPSCs on deep than superficial PNs (sup: 3.2 ± 0.5, deep: 0.9 ± 0.2 Hz; **Fig. 4A**). These data suggest that while inhibitory input levels are similar between PN subclasses, deep PNs receive substantially lower levels of spontaneous excitatory input, or receive much weaker input that falls below the level of recording noise. Therefore, the overall input-output balance between CA3 PN subclasses appears shifted, whereby superficial PNs receive and integrate greater levels of input, yet deep PNs respond more strongly when receiving excitation.

**Fig. 4.**
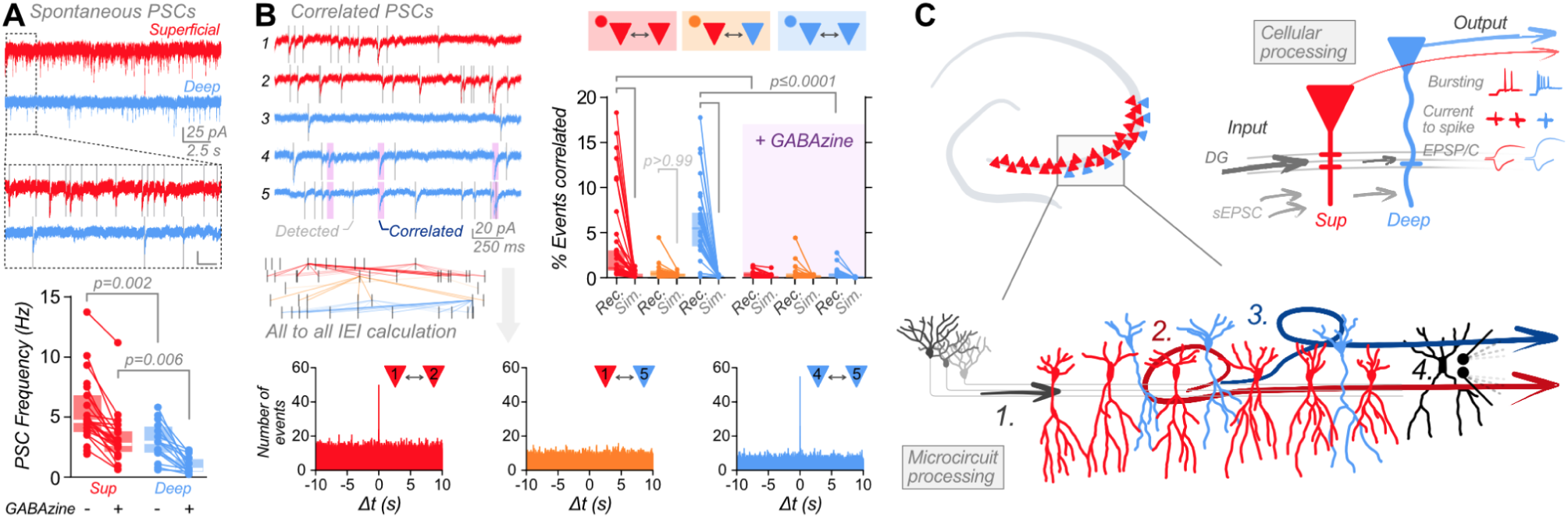
| sPSC timing suggests subclass-dependent inhibitory inputs. **A** sPSC frequency is higher on superficial than deep PNs, suggesting more abundant or stronger synaptic input (sup: 5.37 ± 0.52 Hz, n = 26; deep: 2.96 ± 0.39 Hz, n = 16; Kruskal-Wallis test, p < 0.0001). Bath application of 10 µM GABAzine blocks a similar frequency of events from both subclasses (sup: 2.2 Hz, deep: 2.0 Hz blocked), meaning far less sEPSCs are recorded on deep PNs (sup: 3.20 ± 0.48 Hz, n = 21; deep: 0.94 ± 0.17 Hz, n = 16; Kruskal-Wallis test, p < 0.0001). **B** Detected sPSCs (grey bars) frequently displayed synchronous occurrences on different cells during multicellular recordings (pink highlights). Calculation of all IEIs between event times of different cells, allows plotting of sPSC event correlation between cells (lower). Histograms of IEIs (1 ms bin width) show sharp peaks specifically at Δt = 0 s, demonstrating strictly coincident events between cell pairs. Peaks are observed between superficial cells (Cell 1 vs. 2, red) and between deep cells (Cell 4 vs. 5, blue), but not across classes (Cell 1 vs. 5, orange). Quantification of the percentage of all events that are coincident between cell pairs (Rec. <0.5 ms IEI) shows significant correlation above simulated random levels (Sim.) for within cell class, but not between cell class comparisons (Rec. = recorded, Sim. = simulated. sup-sup (red): Rec – 3.38 ± 0.69%, Sim – 0.24 ± 0.05%, n = 44 pairs; sup-deep (orange): Rec – 0.62 ± 0.15%, Sim – 0.30 ± 0.04%, n = 30 pairs; deep-deep (blue): Rec – 5.92 ± 0.69%, Sim – 0.09 ± 0.02%, n = 34 pairs. Kruskal-Wallis test, p < 0.0001). Bath application of GABAzine (purple) substantially reduces the percentage correlation (sup-sup: Rec – 0.39 ± 0.07%, Sim – 0.10 ± 0.04%, n = 34 pairs; sup-deep: Rec – 0.46 ± 0.16%, Sim – 0.13 ± 0.03%, n = 30 pairs; deep-deep: Rec – 0.43 ± 0.10%, Sim – 0.00 ± 0.00%, n = 32 pairs. Kruskal-Wallis test, p < 0.0001), suggesting inhibitory input as a source for these inputs. Data from 12 multicellular recordings. **C** Hippocampal CA3 contains distinct PN subclasses in two layers (deep (blue) and superficial (red), upper left). Intrinsic neuronal properties display altered input output balance, with deep neurons receiving less input, but tuned to provide more output when input is received (upper right). These subclasses form a two-layer processor in the CA3 recurrent circuit (lower), potentially allowing parallel computations in the CA3 network. GC input (1.) is predominantly received by superficial PNs, which form a recurrent network (2.) and project both downstream and into the second recurrent network of deep PNs (3.). Specific innervation of each PN subclass by distinct interneurons allows gating of information flow through each sublayer (black, 4.).

Through recording multiple cells simultaneously, we observed instances where sPSCs occurred synchronously on two different cells. To determine whether this observation occurred by coincidence, or through circuit mechanisms, we detected sPSCs and computed matrices of interevent intervals (IEIs) between all events on all recorded pairs of cells (**Fig. 4B** see also (Rieubland et al., 2014)). Plotting event crosscorrelograms with fine, 1-ms time bins showed striking peaks at Δt = 0 for a subpopulation of cell pairs. This indicated that a proportion of exquisitely synchronous inputs occurred well above randomness. As these events were not shared across all recorded cells, they appear to result from activation of an unrecorded neuron presynaptic to two recorded neurons. Therefore the timing of sPSCs can be used to infer the wiring of inputs onto recorded cells. Quantifying the percentage of events that are correlated between each cell pair revealed substantial shared input onto pairs of both superficial and deep PNs (3.4 ± 0.7% and 5.9 ± 0.7% respectively). Simulation of random event timings shows that the correlated input levels we observed were well above what would be expected by chance. Surprisingly however, we did not observe these coincident, shared inputs onto cells of different classes (0.6 ± 0.2% synchronous, **Fig. 4B**). These results suggest that a presynaptic population of neurons specifically wires into each PN subclass. Application of GABAzine substantially reduced the percentage of correlated events (**Fig. 4B**). Therefore, CA3 PN subclasses receive inhibition from distinct populations of interneurons, which may allow layer-specific control of recurrent circuit function.

## Discussion

Our study reveals several unexpected properties of the hippocampal CA3 circuit. First, our results shed new light on the heterogeneity of principal neurons in CA3. Multiple studies have highlighted CA3 PN diversity on the deep-superficial axis. These include functional differences, with bursting and non-bursting cells (Masukawa et al., 1982; Bilkey and Schwartzkroin, 1990; Hemond et al., 2008; Raus Balind et al., 2019), but also morphological differences, such as long– and short-shaft PNs with differing mossy fiber input levels (Fitch et al., 1989) or the timing of neuronal generation (Marissal et al., 2012). Consistent with recent results (Magó et al., 2021), we demonstrate that deep neurons are a distinct class of CA3 PNs, with burst firing phenotype and sparse, but not absent mossy-fiber input. This deep PN class includes morphologically identified ‘athorny’ neurons (Hunt et al., 2018), but we find that complete lack of thorny excrescences is not a strict requirement for subclass identity. Genetic determination of how this ‘deep’ subclass maps onto previously identified genetic divisions in CA3 sublayers (Thompson et al., 2008; Yao et al., 2021), or how sparse PCP4+ cells in CA3 relate to this population (Fernandez-Lamo et al., 2019) is required for complete subclass characterization. Deep and superficial layer organization is reminiscent of CA1, in which superficial and deep cells can be distinguished based on morphological properties, calbindin expression, and synaptic connectivity (Lorente de Nó, 1934; Celio, 1990; Mizuseki et al., 2011; Navas-Olive et al., 2020; Soltesz and Losonczy, 2018; Danielson et al., 2016; Morris et al., 1995; Lee et al., 2014). Thus, two-layer organization is a general principle that applies throughout the hippocampal formation.

Our results reveal striking asymmetry in hippocampal connectivity. CA3 is not a single recurrent network as previously assumed. Superficial and deep cells form intralayer recurrent networks, with almost unidirectional activity flow from superficial to deep PNs, at least at the local level. Accordingly, our view of the hippocampal trisynaptic circuit needs revision. While superficial CA3 PNs are embedded in the classical ‘trisynaptic circuit’, forming an EC→GC→sup-CA3→CA1 multisynaptic loop, low DG input and asymmetric connectivity of deep CA3 cells forms a nested ‘tetrasynaptic’ subcircuit: EC→GC→sup-CA3→deep-CA3→CA1 (**Fig. 4C**). In addition, how maintenance of the laminar output of CA3 follows into deep and superficial CA1 layers will also have consequences for hippocampal processing, and must be explored in the future.

The potential processing power of the microcircuit arrangement we observe is an interesting consideration, as highlighted by the current utility of multilayer recurrent networks for complex, artificial intelligence operations. Ensembles of PNs in the CA3 recurrent system are thought to build and store our experiences (Treves and Rolls, 1994). Given that the deep PN layer will receive the output of these ‘completed’ ensembles, their potential for higher-order associations will be interesting to study. In addition, the altered input-output balance of deep and superficial PNs (**Fig. 4C**) will mean ensemble size and activation threshold will likely be different between layers, further complexifying the processing power. One caveat of multicellular patch clamp-based circuit mapping is the focus on local connectivity. It is possible that deep to superficial connectivity exists, but on a long distance scale. Such an arrangement could broadcast local CA3 dynamics for long-range coordination of network events. Future work, for example using transsynaptic labeling with rabies viruses (Sumser et al., 2022), will be needed to map the synaptic inputs and outputs of the local, two layered CA3 processor that we have identified. Previous work has suggested a role for deep (athorny) PNs in coordinating network events, such as sharp-wave ripples (Hunt et al., 2018). Our asymmetric connectivity measurements suggest that this action does not occur through local recurrent activity of deep PNs, however both long-range connectivity or local disinhibitory mechanisms are possible.

The hippocampus is part of the allocortex, a phylogenetically old and highly conserved brain region. It is generally thought that the organization of the allocortex with its 3-layered structure and its linear connectivity scheme is much simpler than that of the neocortex with its 6 layers and several overlapping canonical connectivity motifs (Douglas and Martin, 2004). Our results suggest that the hippocampal CA3 network and the canonical neocortical circuit might be more similar to each other than previously thought. Not only the presence of two layers throughout the hippocampal trisynaptic circuit, but also the preferential connections from superficial to deep CA3 PNs are reminiscent of the neocortex (Douglas and Martin, 2004). Biased connectivity towards deeper neurons has been observed in both mouse subiculum (Peng et al., 2017), and within human neocortical layers (Peng et al., 2024). Thus, while somatic density has been condensed in the allocortex, the circuit architectures of allo– and neocortex may be more similar than appreciated.

Previous research has demonstrated that deep PNs are the first to be generated in hippocampal development (Marissal et al., 2012), while in CA1, neurons generated at the same developmental time point have been suggested to be preferentially connected to the same interneuron populations (Huszár et al., 2022). Our results are in line with these observations, indicating that deep and superficial PNs share innervation from distinct interneurons. The topology of this inhibitory wiring has the potential to gate activity flow between layers. Therefore, identifying these interneuron populations and determining their complete integration into the CA3 microcircuit will further illuminate the complexity of the CA3 processor.

The hippocampal CA3 network is the largest autoassociative network in the brain, and plays a key role in memory. Furthermore, the CA3 network is thought to be critically important for higher-order computations, such as pattern separation and pattern completion (Nakashiba et al., 2008; Rolls, 2018). How these two antagonistic functions are employed and integrated within a single circuit remains unknown (Guzman et al., 2021). Previous *in vivo* recordings suggested division of labor, with proximal CA3 being more dedicated to pattern separation, and distal CA3 more specialized on pattern completion (Lee et al., 2015). The shift in cell type abundance, properties, and potentially microcircuit connectivity across CA3 may play a role in the transformation from pattern separation to pattern completion along the transverse axis and regulate the balance between these two computations. Together, our results help to break down the view of a ‘simple’ hippocampal circuit, and begin to identify the microcircuit details which are necessary to understand the complex neural processing that our memory requires.

## Methods

### Animals

All procedures were performed in strict accordance with institutional, national, and European guidelines for animal experimentation, approved by the Bundesministerium für Bildung, Wissenschaft und Forschung of Austria. Wild-type C57BL/6J mice (RRID:IMSR_JAX:000664) or GAD67-EGFP mice (provided by K. Obata, see (Tamamaki et al., 2003; Hosp et al., 2014)) of both sexes were used at postnatal (P) day 20–30 (median age: P23). All animals were housed with *ad libitum* access to food and water, with constant temperature (22°C) and humidity (50–60%), under a 12 hr light-dark cycle, and used for experiments during the light phase.

### Acute slice preparation

Animals were sacrificed by decapitation under light isofluorane anesthesia. Brains were extracted rapidly in ice-cold high-sucrose artificial cerebrospinal fluid (aCSF, either: 64 mM NaCl, 25 mM NaHCO_3_, 2.5 mM KCl, 1.25 mM NaH_2_PO_4_, 10 mM D-glucose, 120 mM sucrose, 7 mM MgCl_2_, and 0.5 mM CaCl at osmolarity ∼334 mOsm; or 87 mM NaCl, 25 mM NaHCO_3_, 2.5 mM KCl, 1.25 mM NaH_2_PO_4_, 10 mM D-glucose, 75 mM sucrose, 7 mM MgCl_2_, and 0.5 mM CaCl at osmolarity ∼325 mOsm) with 95% O_2_ and 5% CO_2_ gas mixture (carbogen). Hemispheres were separated by a single sagittal cut using a scalpel blade. Acute slices were cut using transverse or quasi-transverse preparation (Bischofberger et al., 2006) and glued to a mounting block using liquid superglue (UHU, 45570). 350-µm-thick slices were cut with a VT 1200 vibratome (Leica Microsystems). Slices were then transferred to a recovery chamber containing either high-sucrose aCSF, or recording aCSF (containing 125 mM NaCl, 25 mM NaHCO_3_, 2.5 mM KCl, 1.25 mM NaH_2_PO_4_, 25 mM D-glucose, 2 mM CaCl_2_, and 1 mM MgCl_2_, osmolarity ∼317 mOsm) at 35°C for 30–45 mins with continuous carbogen bubbling, before maintenance in this solution at room temperature (RT, 20–22°C) until recording.

### Electrophysiology

Slices were transferred to the recording chamber, and continuously perfused with aCSF bubbled with carbogen. Slices were held in place with a platinum harp with nylon threads to prevent tissue movement during recordings. Patch-clamp recording pipettes were pulled from thick-walled borosilicate glass tubing (Hilgenberg, 2 mm OD, 1 mm ID, 1807542), and filled with intracellular solution (containing 135 mM K gluconate, 20 mM KCl, 0.1 mM EGTA, 2 mM MgCl_2_, 2 mM Na_2_ATP, 0.3 mM NaGTP, and 10 mM HEPES, adjusted to pH 7.28 with KOH; osmolarity ∼302 mOsm, with 0.2% (w/v) biocytin). Micropipettes had open tip resistances of 2–6 MΩ when filled with internal solution, and were positioned manually using eight Junior 20ZR micromanipulators (Luigs and Neumann). Neurons were targeted using infrared differential interference contrast (IR-DIC) videomicroscopy based on their soma location in the CA3 pyramidal cell layer. Electrical signals were recorded using four Multiclamp 700B amplifiers (Molecular Devices), low-pass filtered at 6–10 kHz with built-in Bessel filters, and digitized at 20 kHz with a Power 1401 data acquisition interface (Cambridge Electronic Design). Protocols were generated and applied using Signal 6.0 software (CED). All experiments were performed at 21°C (range: 20–22°C).

Pipette offset and capacitance were measured and accounted for during all recordings. During current-clamp recordings, pipette capacitance was ∼70% compensated in all cases, and series resistance compensation was applied as appropriate. Neuronal firing properties were assessed by injection of 1-s hyperpolarizing or depolarizing currents in 50 pA steps (−100 to 400 pA range). Synaptic connectivity was tested by eliciting action potentials (APs) in each recorded neuron in current-clamp mode in turn, while recording responses from all other neurons in either current– or voltage-clamp configurations. APs were elicited by brief current injection (2–5 ms) at a minimal level to reliably evoke single spikes (typically 1–2 nA). Connectivity was routinely tested by stimulation with 5 APs at 20 Hz, repeated at least 40 times. Monosynaptic connections were typically identified by reliable appearance of synaptic responses in the average trace across all pulses of 20 Hz stimulation, with a short latency between presynaptic spike peak and response onset (< 4 ms). Electrical coupling was tested by injection of a 50 pA hyperpolarizing current for 250 ms to the presynaptic cell, and monitoring responses in all other cells. No electrical coupling was observed between CA3 PNs. After recordings, pipettes were slowly retracted from the cell somata to form outside-out patches, which ensured retention of intracellular biocytin for post-hoc staining. The quality of patch formation and the physical location of recorded cells in the tissue slice were documented for later cell identification in stained tissue. Recorded traces were analyzed using Stimfit (version v0.15.8; (Guzman et al., 2014)) or custom Matlab scripts. Cells with a membrane potential above −50 mV or requiring current injection to maintain V_m_ below −50 mV were excluded from analysis of passive and active properties. Cell properties were tested immediately after achieving the whole-cell configuration. Resting membrane potential was measured as the median V_m_ without current injection (0 pA injection step of firing characterization), input resistance was calculated from the end of −100, −50, and +50 pA injections (median voltages between 0.65 and 0.95 s of 1 s pulses). Rheobase current is the required injection for first spike generation during 50 pA step injections. Maximum bursting frequency was defined as the highest instantaneous frequency of the first two spikes upon current injection at either 350 or 400 pA steps. Initial interspike interval (ISI) is defined as the maximal ISI from the first two sweeps generating at least two spikes. Spike-frequency adaptation was measured from the first sweep with at least five APs calculated by dividing the mean of the first two ISIs by the mean of the last two ISIs (1 therefore represents non-adapting spike frequencies, and values closer to 0 are more adapting). All voltage-clamp recordings were performed with a holding potential of −70 mV. Evoked synaptic successes were defined as sweeps with responses of consistent latency exceeding 2.5 σ (EPSCs) or 3 σ (EPSPs) of baseline noise immediately prior to AP generation. Synaptic current rise times are between 20–80% of maximal response, while decay kinetics are time constant of a monoexponential fit. Voltage-clamp synaptic responses were only analyzed in pairs with a postsynaptic access resistance of less than 20 MΩ.

sPSCs were recorded for at least 5 min, monitoring voltage-clamp conditions with a test pulse at least every 20 s. Cells that had an access resistance above 20 MΩ were discarded. Events were detected using a template based analysis (minidet.m, Biosig toolbox, http://biosig.sf.net/; see (Jonas et al., 1993; Clements and Bekkers, 1997)), and event times were measured as event onset. Crosscorrelograms were plotted by calculation of IEIs between all detected events of each cell, and all detected events of other recorded cells. ‘Correlated events’ were defined as events with onset times within 0.5 ms of one another (correlation window = event onset ± 0.5 ms). The percentage of correlated events between each pair of recorded neurons was calculated for both cells of the pair. To confirm correlated event observation was not a result of by-chance superposition, the expected random correlated event frequency was simulated by randomisation of event times for each cell and recalculation of correlated event percentages for each pair. sEPSCs were pharmacologically isolated by application of 10 μM GABAzine to aCSF.

A subset of recorded CA3 PNs from this dataset (1272 tested connections) was previously included in a manuscript for cross species connectivity comparison (Watson et al., 2024).

### Post-hoc morphological visualization

After recording, all slices were fixed and stained for intracellular biocytin to assess neuronal morphology. Slices were routinely imaged using 3,3’-diaminobenzidine (DAB) as chromogen; however a subset of slices were imaged using fluorescence-labeling. For DAB staining (Vectastain ABC-Elite Standard kit, Vector Laboratories PK-6100), slices were fixed in a solution containing 2.5% paraformaldehyde (PFA, TAAB Laboratories Equipment), 1.25% glutaraldehyde (GA; CarlRoth, 4157.1), and 15% (v/v) saturated picric acid solution (Sigma-Aldrich, P6744-1GA). After washing with phosphate buffer (PB; 0.1 M NaH_2_PO_4_ and 0.1 M Na_2_HPO_4_, Merck; titrated to pH 7.35), samples were treated with hydrogen peroxide (1%, 10 min; Sigma-Aldrich, 95321-100ml), permeabilized with 2% Triton X-100 (in PB, Sigma-Aldrich) for 1 h, and transferred to a solution containing 1% avidin-conjugated horseradish peroxidase complex and 1% Triton X-100 for ∼12 h. After rinsing in 0.1 M PB, slices were incubated with a solution containing 0.036% DAB (Sigma-Aldrich, D5637-5G), 0.006% NiCl_2_ (Sigma-Aldrich, 223387-25G) and 0.008% CoCl_2_ (Sigma-Aldrich, C8661-25G), and developed by addition of 0.01% hydrogen peroxide. DAB-developed slices were mounted in Mowiol and cured for at least 24 hours, before visualization using an Olympus BX61 widefield microscope. Recorded cells were visualized using 4, 10, or 20x objectives.

For fluorescent staining, slices were fixed with 4% (w/v) paraformaldehyde (PFA) in 0.1 M PB at 4°C and washed in 0.1 M PB after 24 h to stop the fixation reaction. Slices were blocked and permeabilised by incubation with solution containing 5% natural goat serum (NGS; Biozol ENG9010-10) and 0.4% Triton X-100 in 0.1 M PB for 2–3 hours at RT. Alexa Fluor (AF) 647-conjugated streptavidin (1:300 diluted from 2 mg ml^−1^ stock solution, Invitrogen S32357) was added, and incubated overnight in blocking/permeabilising solution at RT with gentle shaking. Samples were then washed in 0.1 M PB (3 x 30 min), before incubation with DAPI (4’,6-diamidino-2-phenylindole, dilactate, 0.1 µg ml^−1^ final concentration in PB, Invitrogen D3571) in 0.1 M PB for 10–20 mins at RT. Recorded slices were cleared by incubation for 10 min at RT in CUBIC solution (Tainaka et al., 2014) (50% sucrose, 25% urea, 10% 2,2’,2’’,-nitrilotriethanol, 0.1% and Triton X-100 in MilliQ water; all Sigma-Aldrich: 16104, U5128-500G, 90279-100ml), before mounting on glass slides (Assistant, KarlHecht Ref:42406020) beneath a 1.5 thickness glass coverslip (VWR 631-0147) in CUBIC solution surrounded by a ring of Mowiol (Mowiol 4-88, Carl Roth, 713,2; Glycerol, Sigma-Aldrich, G-9012) to prevent exposure of CUBIC solution to air. Samples were cured for at least 24 hours prior to imaging. Slides were imaged on an ANDOR Dragonfly microscope (Oxford Instruments) equipped with a Zyla 4.2 Megapixel sCMOS camera (2048 x 2048 pixels), using either a 10x air objective (Nikon MRD00105, CFI P-Apo 10x, NA 0.45) or a 20x water-immersion objective (Nikon MRD77200, CFI P-Apochromat 20x, NA 0.95). A pinhole disc with 25 µm hole diameters was used. Tiles were stitched using Imaris Stitcher software (Oxford Instruments).

Intersomatic distances were calculated as Euclidean distance between somatic centers in three dimensions. Proximal-distal distance dependence to connectivity and somatic location on this axis was measured by manual segmentation of *stratum pyramidale* as a vector to the end of *stratum lucidum* (from 4x images), and determination of somatic location along this vector. Somatic depth was taken as the distance between somatic center and the upper bound of *stratum lucidum*.

### Cell Classification

CA3 PN classification into ‘deep’ or ‘superficial’ subclasses was performed using a multi-step procedure. First, cells were provisionally classified morphologically as classical ‘thorny’ PNs, or PNs with athorny cellular morphology (deep somatic location and thin primary dendrite with long distance to first branch point (typically over 50 μm), following (Hunt et al., 2018)), either without or with visible thorny excrescences. Next we performed LDA classification from depth/bursting scatter plots of only these morphologically identified neurons with perfect recording conditions (resting membrane potential < −50 mV, no current injection required to maintain V_mem_, R_input_ in the range 20–500 MΩ). Requirements for ‘deep’ PN classification were then set as, somatic depth-burst firing relationship beyond the LDA segregation cut off ([Max burst freq, Hz] = −1.29 × [Somatic depth, μm] + 124.37, as well as passing minimal cutoffs of somatic depth > 30 μm, maximum burst frequency > 40 Hz, and initial ISI of < 0.1 s. All athorny-like neurons (with or without thorns) in the ‘deep-bursting’ cluster by this definition, were classified as ‘deep’ PNs, while all classical thorny neurons outside these criteria were classified as superficial neurons. 16 cells had a mismatch between clustering based classification and provisional morphological classification. These cells were morphologically reassessed and 8 were assigned to their respective LDA-based classification. 2 apparent ‘superficial athorny’ cells, and 3 PNs with deep somatic location but immediate dendritic branching and abundant thorny excrescences, were deemed unclassifiable and were excluded. These criteria were extended to classify 52 neurons which had incomplete morphology but measurable somatic depth and adequate firing properties. Cells without measurable somatic depth or suboptimal recording quality were excluded from further analysis (n = 164).

### Statistical analysis

All data are reported as mean ± SEM in text and figure legends unless otherwise specified. Box plots depict data as median (line), 25th and 75th quartile (box) and min/max points (whiskers). Symbols with errors depict mean ± SEM (Fig. 3C). Connectivities are presented as bars (measured data) with standard deviation estimated from a binomial distribution. Statistical tests are reported in figure legends in all cases. As biological datasets rarely exhibit a normal distribution, non-parametric statistical tests for either two-sample (Mann-Whitney test) or multi-sample (Kruskal-Wallis test) data were applied as appropriate, unless specified otherwise. Where Kruskal-Wallis test has been applied, pairwise comparisons presented on graphs are a result of Dunn’s multiple comparison test. Statistical tests were performed using GraphPad Prism 10, and exact p values were reported throughout the manuscript. Proportional datasets (e.g. for connectivity) were compared using Fisher’s exact test, either for multisample data (employed in R), or as individual pairwise comparisons (run using GraphPad Prism) with a Benjamini-Hochberg correction for multiple comparisons.

## Acknowledgements

We thank Andrea Navas-Olive and Rebecca J. Morse-Mora for critically reading an earlier version of the manuscript. We also thank Florian Marr and Christina Altmutter for excellent technical assistance, Alois Schlögl for programming and data-handling assistance, Todor Asenov for technical support, and Eleftheria Kralli-Beller for manuscript editing. This research was supported by the Scientific Services Units (SSUs) of ISTA. We are particularly grateful for assistance from the Imaging and Optics Facility, Preclinical Facility, Life Science Facility, and Miba Machine Shop. The project received funding from the European Research Council (ERC) under the European Union’s Horizon 2020 research and innovation programme (grant agreement No 692692 to P.J.; Marie Skłodowska-Curie Actions Individual Fellowship No. 101026635 to J.F.W.) and the Austrian Science Fund (PAT 4178023 to P.J.).

## Author Contributions

J.F.W. and P.J. conceived the project, J.F.W. performed experiments and analyzed data, V.V.-B. supported electrophysiology experiments and analysis, J.F.W. and P.J. wrote the paper and acquired funding for the project. All authors jointly revised the paper.

## Supplementary Information

**Supplementary Fig. 1.**
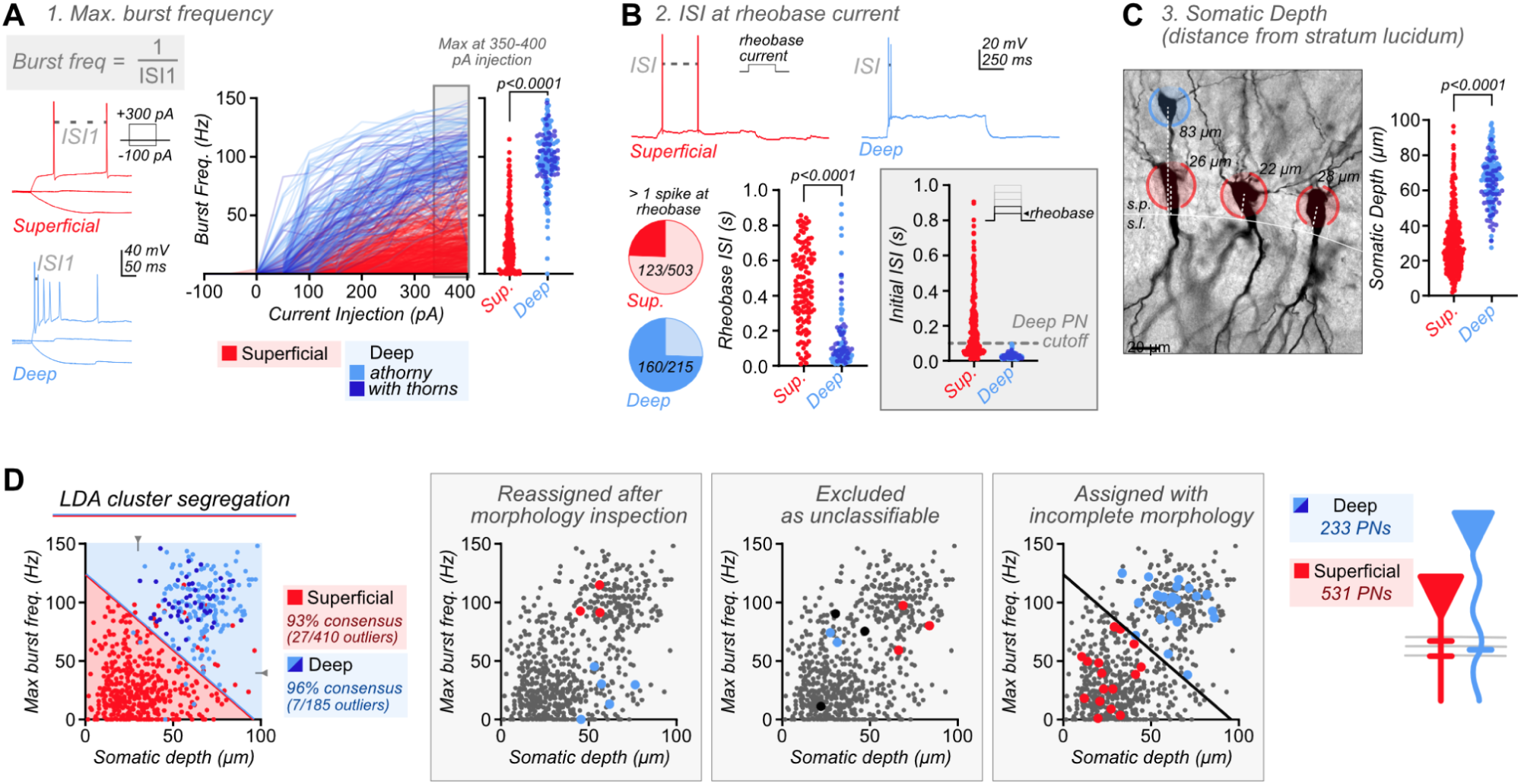
| Categorisation of neurons into deep and superficial subclasses. Three parameters were used to define CA3 PN subclasses: maximal frequency of burst firing upon current injection (**A**), burst firing frequency around rheobase current (**B**), and somatic depth in *stratum pyramidale* (s.p., **C**). **A** Burst frequency on increasing current injection (1/ first ISI) shows distinct trajectories for morphologically identified ‘superficial/thorny’ (red) and ‘deep/athorny’ neurons (blue). Sparsely thorny neurons with athorny morphology (deep, with thorns, dark blue) follow the athorny trajectory. Max. frequency is measured as the maximum frequency at 350 or 400 pA current injection (gray box, sup: 27.9 ± 1.1 Hz, n = 496; deep (athorny): 97.3 ± 2.2 Hz, n = 156; deep (with thorns): 100.9 ± 2.5 Hz, n = 49; Kruskal-Wallis test, p < 0.0001). **B** During 1-s depolarisations at rheobase current, the majority of deep neurons fire more than 1 AP, while this is much rarer for superficial neurons (sup: 123/503; deep: 160/215; Fisher’s exact test, p < 0.0001). Of these occurrences, the first ISI is distributed across the entire 1-s window for superficial neurons, yet is systematically fast for deep neurons (sup: 0.45 ± 0.02 s, n = 123; deep (athorny): 0.13 ± 0.02 s, n = 122; deep (with thorns): 0.14 ± 0.02; Kruskal-Wallis test, p < 0.0001). Calculating the maximal ISI from the first two sweeps giving at least 2 spikes (initial ISI, gray box), allowed definition of a cutoff for consideration as a deep PN (Initial ISI < 0.1 s) (sup: 0.20 ± 0.01 s, n = 449; deep (athorny): 0.03 ± 0.00 s, n = 164; deep (with thorns): 0.03 ± 0.00 s, n = 50; Kruskal-Wallis test, p < 0.0001). **C** Somatic depth, the Euclidean distance between somatic center and *stratum pyramidale/stratum lucidum* boundary, was significantly greater for putative deep neurons than superficial neurons (sup: 30.6 ± 0.8 μm, n = 503; deep (athorny): 67.7 ± 1.0 μm, n = 165; deep (with thorns): 60.6 ± 1.9 μm, n = 50; Kruskal-Wallis test, p < 0.0001). Multiple comparisons tests for all parameters were significantly different between superficial and deep classes (all p < 0.0001), and no different between deep subgroups (max burst frequency, p > 0.99; Initial ISI, p > 0.99; somatic depth, p = 0.44). **D** LDA subclass identification from scatter plot of maximal burst frequency (from **A**) and somatic depth (from **C**) showed 94% agreement with morphological identification (93% and 96% agreement with superficial and deep classifications, respectively). Equation of linear discriminator: [Max. burst freq., Hz] = –1.29 × [Somatic depth, μm] + 124.37. Minimum depth and burst firing cutoffs were also applied for ‘deep PN’ classification (grey arrows). As deep (with thorns) neurons show no difference to athorny neurons in any parameter, and fall exclusively into the ‘deep’ subcluster, these neurons are considered one indivisible subclass. LDA subclassification allowed reassignment of misidentified neurons (n = 8), and exclusion of PNs which did not fit subclass identities (n = 8). In addition, neurons with recorded somatic depth, but incomplete morphology could be assigned unambiguously to either subclass (see **Methods** for full details of classification). This procedure resulted in 233 putative deep, and 531 putative superficial CA3 PNs in our recorded data.

**Supplementary Fig. 2.**
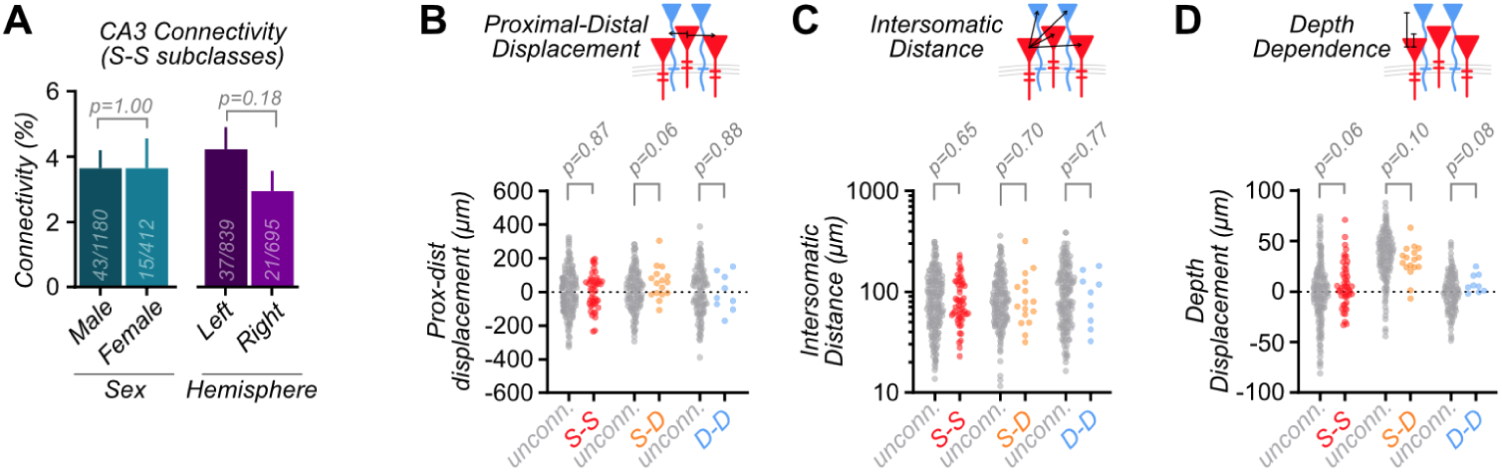
| Spatial dependence of CA3 subclass connectivity. A. No difference in connectivity was observed between male and female mice, or between left and right hemisphere. Data presented is only S-S connectivity (data on bars, Fisher’s exact test, male/ female: p = 1.00; left/ right: p = 0.18). **B** No bias for connectivity in either proximal or distal direction was observed for CA3 PN connections (S-S: unconnected, 0.14 ± 2.9 μm, n = 1485; connected, 1.7 ± 13.8 μm, n = 55; Mann-Whitney test, p = 0.87. S-D: unconnected, 4.4 ± 4.5 μm, n = 518; connected, 56.3 ± 24.5 μm, n = 16; Mann-Whitney test, p = 0.06. D-D: unconnected, 0.6 ± 6.5 μm, n = 375; connected, −4.9 ± 36.6 μm, n = 9; Mann-Whitney test, p = 0.88). **C** No distance dependence to connectivity is observed by comparing intersomatic distances of connected (colored) or unconnected (grey) pairs (S-S: unconnected, 95 ± 1.4 μm, n = 1495; connected, 89.9 ± 6.5 μm, n = 57; Mann-Whitney test, p = 0.65. S-D: unconnected, 96.9 ± 2.3 μm, n = 535; connected, 98.2 ± 17.7 μm, n = 16; Mann-Whitney test, p = 0.70. D-D: unconnected, 110.1 ± 3.6 μm, n = 377; connected, 98.8 ± 17.1 μm, n = 9; Mann-Whitney test, p = 0.77). **D** The dependence of connectivity on intersomatic depth between connected neurons (coloured by subclass) (S-S: unconnected, −0.2 ± 0.6 μm, n = 1516; connected, +6.1 ± 2.7 μm, n = 58; Mann-Whitney test, p = 0.06. S-D: unconnected, 35.9 ± 0.9 μm, n = 540; connected, 29.7 ± 3.9 μm, n = 17; Mann-Whitney test, p = 0.10. D-D: unconnected, −0.4 ± 0.8 μm, n = 385; connected, 7.7 ± 3.1 μm, n = 9; Mann-Whitney test, p = 0.08).

**Supplementary Fig. 3.**
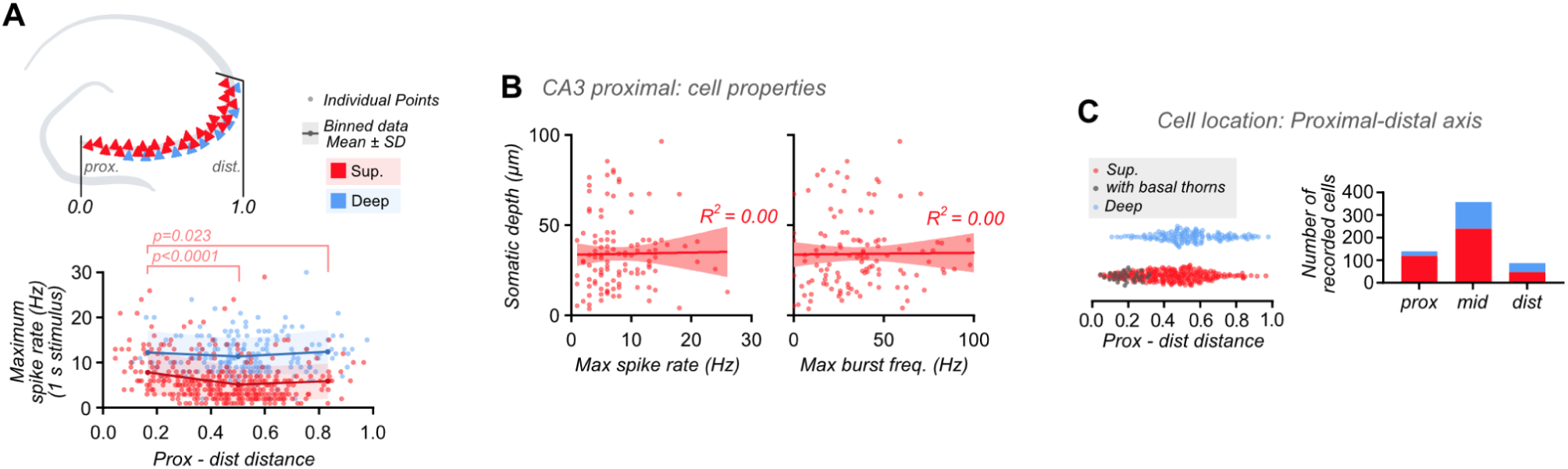
| CA3 firing pattern changes on the proximal-distal axis. A. Scatter plots of maximal firing frequency achieved during 1-s depolarisations (up to 400 pA) show greater frequency in proximal CA3 (close to DG) than distal CA3 (close to CA2). The firing frequency of deep PNs does not appear to be influenced by this axis (blue). Individual data points are plotted with a binned overlay. Binned data is plotted as Mean ± SD (Mean ± SD; sup: proximal – 7.83 ± 4.95 Hz, n = 120; mid – 5.13 ± 3.89 Hz, n = 240; distal: – 5.89 ± 3.86 Hz, n = 46. deep: proximal – 12.25 ± 4.73 Hz, n = 20; mid – 11.37 ± 3.94 Hz, n = 119; distal: – 12.4 ± 4.90 Hz, n = 41. Kruskal-Wallis test, p < 0.0001. Dunn’s multiple comparison test p-values displayed). **B** In proximal CA3 (prox-dist distance < 0.33), there is no correlation between somatic depth and maximal spike rate, or initial burst frequency for superficial PNs. Therefore there is no indication more active proximal PNs include a subpopulation of deep PNs (linear regressions for both parameters show R^2^ < 0.005). **C** The abundance of recorded neurons on the proximal distal axis is biased to mid CA3. Amongst these recorded cells, fewer deep PNs were recorded in proximal CA3. This lower recorded abundance correlates with the presence of the infrapyramidal blade of the mossy fiber tract, which can be detected by the presence of thorny excrescences on basal dendrites (gray) (prox: 120 sup, 20 deep; mid: 240 sup, 119 deep; dist: 46 sup, 41 deep). Note that recorded cell numbers are not a quantitative measure of cell proportions in tissue, due to experimental recording bias.

**Supplementary Fig. 4.**
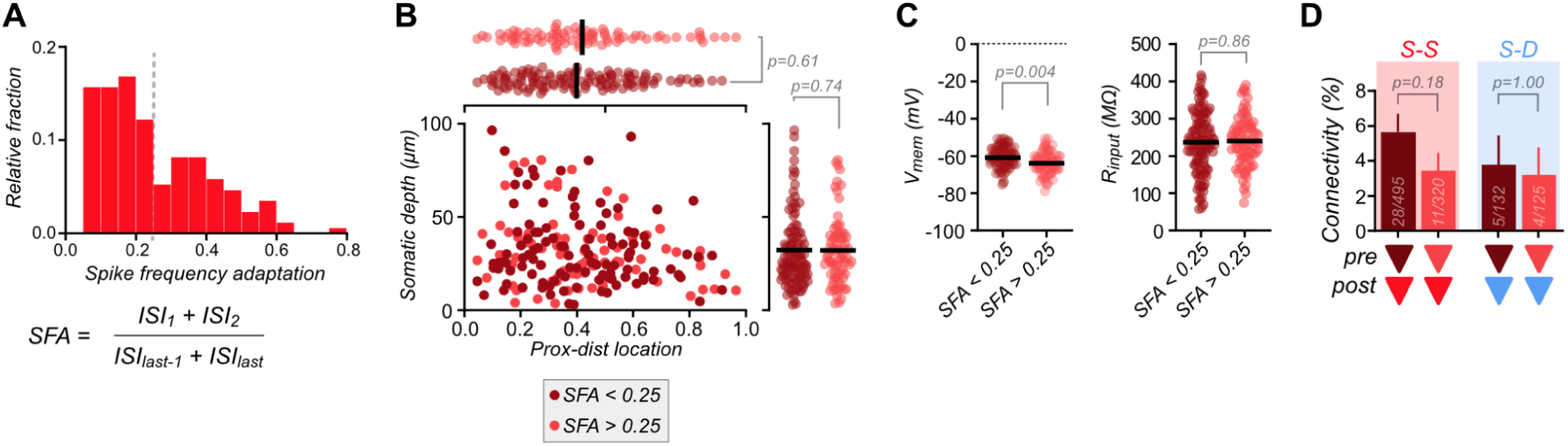
| Superficial neurons with different spike adaptation phenotypes show similar distribution and connectivity. A. Plotting spike frequency adaptation (SFA) for superficial neurons reflects previously reported subgroups (Balleza-Tapia et al., 2022). **B** Superficial neurons with SFA above or below 0.25 show overlapping location on both proximal-distal and deep-superficial axes. Bars indicate mean values. (Depth: ‘SFA<0.25’ – 32.4 ± 1.8 μm, n = 118; ‘SFA>0.25’ – 32.2 ± 2.0 μm, n = 83; Mann-Whitney test, p = 0.74. Prox-dist location: ‘SFA<0.25’ – 0.40 ± 0.02, n = 118; ‘SFA>0.25’ – 0.42 ± 0.02, n = 83. Mann-Whitney test, p = 0.61). **C** Membrane potential is subtly different, but input resistance is unchanged between these subgroups (Vm: ‘SFA<0.25’ – −60.9 ± 0.5 mV, n = 118; ‘SFA>0.25’ – −63.8 ± 0.7 μm, n = 83; Mann-Whitney test, p = 0.004. Input resistance: ‘SFA<0.25’ – 236.4 ± 7.4 MΩ, n = 118; ‘SFA>0.25’ – 239.7 ± 7.4, n = 83. Mann-Whitney test, p = 0.86). **D** Both superficial neuron subgroups show comparable connectivity to both other superficial neurons (data displayed on bars. Fisher’s exact test, p = 0.18), and to deep PNs (Fisher’s exact test, p = 1.00).

## References

1. Amaral, D.G., Witter, M.P., 1989. The three-dimensional organization of the hippocampal formation: A review of anatomical data. Neuroscience 31, 571–591. 10.1016/0306-4522(89)90424-7

2. Andersen, P., Bliss, T.V.P., Skrede, K.K., 1971. Lamellar organization of hippocampal excitatory pathways. Exp. Brain Res. 13, 222–238. 10.1007/BF00234087

3. Balleza-Tapia, H., Arroyo-García, L.E., Isla, A.G., Loera-Valencia, R., Fisahn, A., 2022. Functionally-distinct pyramidal cell subpopulations during gamma oscillations in mouse hippocampal area CA3. Prog. Neurobiol. 210, 102213. 10.1016/j.pneurobio.2021.102213

4. Ben-Simon, Y., Kaefer, K., Velicky, P., Csicsvari, J., Danzl, J.G., Jonas, P., 2022. A direct excitatory projection from entorhinal layer 6b neurons to the hippocampus contributes to spatial coding and memory. Nat. Commun. 13, 4826. 10.1038/s41467-022-32559-8

5. Bilkey, D.K., Schwartzkroin, P.A., 1990. Variation in electrophysiology and morphology of hippocampal CA3 pyramidal cells. Brain Res. 514, 77–83. 10.1016/0006-8993(90)90437-G

6. Bischofberger, J., Engel, D., Li, L., Geiger, J.R.P., Jonas, P., 2006. Patch-clamp recording from mossy fiber terminals in hippocampal slices. Nat. Protoc. 1, 2075–2081. 10.1038/nprot.2006.312

7. Celio, M.R., 1990. Calbindin D-28k and parvalbumin in the rat nervous system. Neuroscience 35, 375–475. 10.1016/0306-4522(90)90091-h

8. Cembrowski, M.S., Spruston, N., 2019. Heterogeneity within classical cell types is the rule: lessons from hippocampal pyramidal neurons. Nat. Rev. Neurosci. 20, 193–204. 10.1038/s41583-019-0125-5

9. Clements, J.D., Bekkers, J.M., 1997. Detection of spontaneous synaptic events with an optimally scaled template. Biophys. J. 73, 220–229. 10.1016/S0006-3495(97)78062-7

10. Danielson, N.B., Zaremba, J.D., Kaifosh, P., Bowler, J., Ladow, M., Losonczy, A., 2016. Sublayer-specific coding dynamics during spatial navigation and learning in hippocampal area CA1. Neuron 91, 652–665. 10.1016/j.neuron.2016.06.020

11. Douglas, R.J., Martin, K.A.C., 2004. Neuronal circuits of the neocortex. Annu. Rev. Neurosci. 27, 419–451. 10.1146/annurev.neuro.27.070203.144152

12. Dudek, S.M., Alexander, G.M., Farris, S., 2016. Rediscovering area CA2: unique properties and functions. Nat. Rev. Neurosci. 17, 89–102. 10.1038/nrn.2015.22

13. Fernandez-Lamo, I., Gomez-Dominguez, D., Sanchez-Aguilera, A., Oliva, A., Morales, A.V., Valero, M., Cid, E., Berenyi, A., Menendez de la Prida, L., 2019. Proximodistal organization of the CA2 hippocampal area. Cell Rep. 26, 1734–1746.e6. 10.1016/j.celrep.2019.01.060

14. Fitch, J.M., Juraska, J.M., Washington, L.W., 1989. The dendritic morphology of pyramidal neurons in the rat hippocampal CA3 area. I. Cell types. Brain Res. 479, 105–114. 10.1016/0006-8993(89)91340-1

15. Guzman, S.J., Schlögl, A., Espinoza, C., Zhang, X., Suter, B.A., Jonas, P., 2021. How connectivity rules and synaptic properties shape the efficacy of pattern separation in the entorhinal cortex–dentate gyrus–CA3 network. Nat. Comput. Sci. 1, 830–842. 10.1038/s43588-021-00157-1

16. Guzman, S.J., Schlögl, A., Frotscher, M., Jonas, P., 2016. Synaptic mechanisms of pattern completion in the hippocampal CA3 network. Science 353, 1117–1123. 10.1126/science.aaf1836

17. Guzman, S.J., Schlögl, A., Schmidt-Hieber, C., 2014. Stimfit: quantifying electrophysiological data with Python. Front. Neuroinformatics 8, 16. 10.3389/fninf.2014.00016

18. Hemond, P., Epstein, D., Boley, A., Migliore, M., Ascoli, G.A., Jaffe, D.B., 2008. Distinct classes of pyramidal cells exhibit mutually exclusive firing patterns in hippocampal area CA3b. Hippocampus 18, 411–424. 10.1002/hipo.20404

19. Hosp, J.A., Strüber, M., Yanagawa, Y., Obata, K., Vida, I., Jonas, P., Bartos, M., 2014. Morpho-physiological criteria divide dentate gyrus interneurons into classes. Hippocampus 24, 189–203. 10.1002/hipo.22214

20. Hunt, D.L., Linaro, D., Si, B., Romani, S., Spruston, N., 2018. A novel pyramidal cell type promotes sharp-wave synchronization in the hippocampus. Nat. Neurosci. 21, 985–995. 10.1038/s41593-018-0172-7

21. Huszár, R., Zhang, Y., Blockus, H., Buzsáki, G., 2022. Preconfigured dynamics in the hippocampus are guided by embryonic birthdate and rate of neurogenesis. Nat. Neurosci. 25, 1201–1212. 10.1038/s41593-022-01138-x

22. Jonas, P., Major, G., Sakmann, B., 1993. Quantal components of unitary EPSCs at the mossy fibre synapse on CA3 pyramidal cells of rat hippocampus. J. Physiol. 472, 615–663. 10.1113/jphysiol.1993.sp019965

23. Kowalski, J., Gan, J., Jonas, P., Pernía-Andrade, A.J., 2016. Intrinsic membrane properties determine hippocampal differential firing pattern in vivo in anesthetized rats. Hippocampus 26, 668–682. 10.1002/hipo.22550

24. Lee, H., Wang, C., Deshmukh, S.S., Knierim, J.J., 2015. Neural population evidence of functional heterogeneity along the CA3 transverse axis: pattern completion versus pattern separation. Neuron 87, 1093–1105. 10.1016/j.neuron.2015.07.012

25. Lee, S.-H., Marchionni, I., Bezaire, M., Varga, C., Danielson, N., Lovett-Barron, M., Losonczy, A., Soltesz, I., 2014. Parvalbumin-positive basket cells differentiate among hippocampal pyramidal cells. Neuron 82, 1129–1144. 10.1016/j.neuron.2014.03.034

26. Linaro, D., Levy, M.J., Hunt, D.L., 2022. Cell type-specific mechanisms of information transfer in data-driven biophysical models of hippocampal CA3 principal neurons. PLOS Comput. Biol. 18, e1010071. 10.1371/journal.pcbi.1010071

27. Lorente de Nó, R., 1934. Studies on the structure of the cerebral cortex. II. Continuation of the study of the ammonic system. J. Für Psychol. Neurol. 46, 113–177.

28. Magó, Á., Kis, N., Lükő, B., Makara, J.K., 2021. Distinct dendritic Ca^2+^ spike forms produce opposing input-output transformations in rat CA3 pyramidal cells. eLife 10, e74493. 10.7554/eLife.74493

29. Marissal, T., Bonifazi, P., Picardo, M.A., Nardou, R., Petit, L.F., Baude, A., Fishell, G.J., Ben-Ari, Y., Cossart, R., 2012. Pioneer glutamatergic cells develop into a morpho-functionally distinct population in the juvenile CA3 hippocampus. Nat. Commun. 3, 1316. 10.1038/ncomms2318

30. Masukawa, L.M., Benardo, L.S., Prince, D.A., 1982. Variations in electrophysiological properties of hippocampal neurons in different subfields. Brain Res. 242, 341–344. 10.1016/0006-8993(82)90320-1

31. Miles, R., Wong, R.K.S., 1986. Excitatory synaptic interactions between CA3 neurones in the guinea-pig hippocampus. J. Physiol. 373, 397–418. 10.1113/jphysiol.1986.sp016055

32. Mizuseki, K., Diba, K., Pastalkova, E., Buzsáki, G., 2011. Hippocampal CA1 pyramidal cells form functionally distinct sublayers. Nat. Neurosci. 14, 1174–1181. 10.1038/nn.2894

33. Morris, M.E., Baimbridge, K.G., El-Beheiry, H., Obrocea, G.V., Rosen, A.S., 1995. Correlation of anoxic neuronal responses and calbindin-D_28k_ localization in stratum pyramidale of rat hippocampus. Hippocampus 5, 25–39. 10.1002/hipo.450050105

34. Nakashiba, T., Young, J.Z., McHugh, T.J., Buhl, D.L., Tonegawa, S., 2008. Transgenic inhibition of synaptic transmission reveals role of CA3 output in hippocampal learning. Science 319, 1260–1264. 10.1126/science.1151120

35. Navas-Olive, A., Valero, M., Jurado-Parras, T., de Salas-Quiroga, A., Averkin, R.G., Gambino, G., Cid, E., de la Prida, L.M., 2020. Multimodal determinants of phase-locked dynamics across deep-superficial hippocampal sublayers during theta oscillations. Nat. Commun. 11, 2217. 10.1038/s41467-020-15840-6

36. Peng, Y., Barreda Tomás, F.J., Klisch, C., Vida, I., Geiger, J.R.P., 2017. Layer-specific organization of local excitatory and inhibitory synaptic connectivity in the rat presubiculum. Cereb. Cortex 27, 2435–2452. 10.1093/cercor/bhx049

37. Peng, Y., Bjelde, A., Aceituno, P.V., Mittermaier, F.X., Planert, H., Grosser, S., Onken, J., Faust, K., Kalbhenn, T., Simon, M., Radbruch, H., Fidzinski, P., Schmitz, D., Alle, H., Holtkamp, M., Vida, I., Grewe, B.F., Geiger, J.R.P., 2024. Directed and acyclic synaptic connectivity in the human layer 2-3 cortical microcircuit. Science 384, 338–343. 10.1126/science.adg8828

38. Raus Balind, S., Magó, Á., Ahmadi, M., Kis, N., Varga-Németh, Z., Lőrincz, A., Makara, J.K., 2019. Diverse synaptic and dendritic mechanisms of complex spike burst generation in hippocampal CA3 pyramidal cells. Nat. Commun. 10, 1859. 10.1038/s41467-019-09767-w

39. Rieubland, S., Roth, A., Häusser, M., 2014. Structured connectivity in cerebellar inhibitory networks. Neuron 81, 913–929. 10.1016/j.neuron.2013.12.029

40. Rolls, E.T., 2018. The storage and recall of memories in the hippocampo-cortical system. Cell Tissue Res. 373, 577–604. 10.1007/s00441-017-2744-3

41. Sammons, R.P., Vezir, M., Moreno-Velasquez, L., Cano, G., Orlando, M., Sievers, M., Grasso, E., Metodieva, V.D., Kempter, R., Schmidt, H., Schmitz, D., 2024. Structure and function of the hippocampal CA3 module. Proc. Natl. Acad. Sci. U. S. A. 121, e2312281120. 10.1073/pnas.2312281120

42. Slomianka, L., Amrein, I., Knuesel, I., Sørensen, J.C., Wolfer, D.P., 2011. Hippocampal pyramidal cells: the reemergence of cortical lamination. Brain Struct. Funct. 216, 301–317. 10.1007/s00429-011-0322-0

43. Soltesz, I., Losonczy, A., 2018. CA1 pyramidal cell diversity enabling parallel information processing in the hippocampus. Nat. Neurosci. 21, 484–493. 10.1038/s41593-018-0118-0

44. Squire, L.R., 2009. The legacy of patient H.M. for neuroscience. Neuron 61, 6–9. 10.1016/j.neuron.2008.12.023

45. Squire, L.R., Stark, C.E.L., Clark, R.E., 2004. The medial temporal lobe. Annu. Rev. Neurosci. 27, 279–306. 10.1146/annurev.neuro.27.070203.144130

46. Sumser, A., Joesch, M., Jonas, P., Ben-Simon, Y., 2022. Fast, high-throughput production of improved rabies viral vectors for specific, efficient and versatile transsynaptic retrograde labeling. eLife 11, e79848. 10.7554/eLife.79848

47. Sun, Q., Sotayo, A., Cazzulino, A.S., Snyder, A.M., Denny, C.A., Siegelbaum, S.A., 2017. Proximodistal heterogeneity of hippocampal CA3 pyramidal neuron intrinsic properties, connectivity, and reactivation during memory recall. Neuron 95, 656–672.e3. 10.1016/j.neuron.2017.07.012

48. Tainaka, K., Kubota, S.I., Suyama, T.Q., Susaki, E.A., Perrin, D., Ukai-Tadenuma, M., Ukai, H., Ueda, H.R., 2014. Whole-body imaging with single-cell resolution by tissue decolorization. Cell 159, 911–924. 10.1016/j.cell.2014.10.034

49. Tamamaki, N., Yanagawa, Y., Tomioka, R., Miyazaki, J.-I., Obata, K., Kaneko, T., 2003. Green fluorescent protein expression and colocalization with calretinin, parvalbumin, and somatostatin in the GAD67-GFP knock-in mouse. J. Comp. Neurol. 467, 60–79. 10.1002/cne.10905

50. Thompson, C.L., Pathak, S.D., Jeromin, A., Ng, L.L., MacPherson, C.R., Mortrud, M.T., Cusick, A., Riley, Z.L., Sunkin, S.M., Bernard, A., Puchalski, R.B., Gage, F.H., Jones, A.R., Bajic, V.B., Hawrylycz, M.J., Lein, E.S., 2008. Genomic anatomy of the hippocampus. Neuron 60, 1010–1021. 10.1016/j.neuron.2008.12.008

51. Treves, A., Rolls, E.T., 1994. Computational analysis of the role of the hippocampus in memory. Hippocampus 4, 374–391. 10.1002/hipo.450040319

52. Valero, M., Cid, E., Averkin, R.G., Aguilar, J., Sanchez-Aguilera, A., Viney, T.J., Gomez-Dominguez, D., Bellistri, E., de la Prida, L.M., 2015. Determinants of different deep and superficial CA1 pyramidal cell dynamics during sharp-wave ripples. Nat. Neurosci. 18, 1281–1290. 10.1038/nn.4074

53. Vandael, D., Jonas, P., 2024. Structure, biophysics, and circuit function of a “giant” cortical presynaptic terminal. Science 383, eadg6757. 10.1126/science.adg6757

54. Watson, J.F., Vargas-Barroso, V., Morse-Mora, R.J., Navas-Olive, A., Tavakoli, M.R., Danzl, J.G., Tomschik, M., Rössler, K., Jonas, P., 2024. Human hippocampal CA3 uses specific functional connectivity rules for efficient associative memory. bioRxiv 2024.05.02.592169. 10.1101/2024.05.02.592169

55. Watson, J.F., Vargas-Barroso, V.M., Jonas, P., 2022. Cell-specific synaptic wiring within the hippocampal CA3 network. Soc. Neurosci. Abstr. 019.06.

56. Witter, M.P., 2007. Intrinsic and extrinsic wiring of CA3: Indications for connectional heterogeneity. Learn. Mem. 14, 705–713. 10.1101/lm.725207

57. Yao, Z., van Velthoven, C.T.J., Nguyen, T.N., Goldy, J., Sedeno-Cortes, A.E., Baftizadeh, F., Bertagnolli, D., Casper, T., Chiang, M., Crichton, K., Ding, S.-L., Fong, O., Garren, E., Glandon, A., Gouwens, N.W., Gray, J., Graybuck, L.T., Hawrylycz, M.J., Hirschstein, D., Kroll, M., Lathia, K., Lee, C., Levi, B., McMillen, D., Mok, S., Pham, T., Ren, Q., Rimorin, C., Shapovalova, N., Sulc, J., Sunkin, S.M., Tieu, M., Torkelson, A., Tung, H., Ward, K., Dee, N., Smith, K.A., Tasic, B., Zeng, H., 2021. A taxonomy of transcriptomic cell types across the isocortex and hippocampal formation. Cell 184, 3222–3241.e26. 10.1016/j.cell.2021.04.021

58. Yeckel, M.F., Berger, T.W., 1990. Feedforward excitation of the hippocampus by afferents from the entorhinal cortex: redefinition of the role of the trisynaptic pathway. Proc. Natl. Acad. Sci. U. S. A. 87, 5832–5836. 10.1073/pnas.87.15.5832

